# Function of a novel nasal protrusion for oral-shelling within an adaptive radiation of pupfishes

**DOI:** 10.1101/2020.03.23.004416

**Authors:** Michelle E. St. John, Kristi Dixon, Christopher H. Martin

**Affiliations:** Department of Integrative Biology and Museum of Vertebrate Zoology, University of California, Berkeley, CA 94720, USA; Department of Biology, University of North Carolina at Chapel Hill, 120 South Rd, NC 27599, USA

**Keywords:** adaptive radiation, speciation, novelty, performance, durophagy, craniofacial, foraging

## Abstract

Dietary specialization on hard prey items, such as mollusks and crustaceans, is commonly observed in a diverse array of fish species. Many fish consume these types of prey by crushing the shell to consume the soft tissue within, but a few fishes extricate the soft tissue without breaking the shell using a method known as oral shelling. Oral shelling involves pulling a mollusk from its shell and may be a way to subvert an otherwise insurmountable shell defense. However, the biomechanical requirements and potential adaptations for oral shelling are unknown. Here, we test the hypothesis that a novel nasal protrusion is an adaptation for oral shelling in a durophagous pupfish (*Cyprinodon brontotheroides*). We first demonstrate oral shelling in this species and then predicted that a larger nasal protrusion would allow pupfish to consume larger snails. Durophagous pupfish are found within an endemic radiation of pupfish on San Salvador Island, Bahamas. We took advantage of closely related sympatric species and outgroups to test: 1) whether durophagous pupfish shell and consume more snails than other species, 2) if F1 and F2 durophagous hybrids consume similar amounts of snails as purebred durophagous pupfish, and 3) to determine if nasal protrusion size in parental and hybrid populations increases the maximum diameter snail consumed. We found that durophagous pupfish and their hybrids consumed the most snails, but did not find a strong association between nasal protrusion size and maximum snail size consumed within the parental or F2 hybrid population, suggesting that the size of their novel nasal protrusion does not provide a major benefit in oral shelling. Instead, we suggest that nasal protrusion may increase feeding efficiency, act as a sensory organ, or is a sexually selected trait, and that a strong feeding preference may be most important for oral shelling.

**Significance Statement:** Specialization on hard-shell prey items (i.e. durophagy) is a common dietary niche among fishes. Oral shelling is a rare technique used by some durophagous fish to consume prey items like snails; however, adaptations for oral shelling are still unknown. Here, we document the first evidence of oral shelling in a cyprinodontiform fish, the durophagous pupfish (*Cyprinodon brontotheroides*), and experimentally test whether its novel nasal protrusion is an adaptation for oral shelling using hybrid feeding trials.

## Introduction

Dietary specialization is thought to be one way to reduce competition for a food source or to forage more optimally (Pyke 1984; Futuyman and Moreno 1988; Robinson and Wilson 1998). One form of dietary specialization, especially among fishes, is the increased consumption of hard-shelled prey items, such as mollusks and crustaceans (hereafter referred to as durophagy), and both freshwater and marine fishes include durophagous specialists. There are two main ways that fish consume hard-shelled prey items: First, fish may crush or break the outer shell to consume the soft tissue within. Some fishes, such as black carp (*Mylopharyngodon picesus*), pumpkinseed sunfish (*Lepomis gibbosus*), redear sunfish (*Lepomis microlophus*), black drum (*Pogonias cromis*), Florida pompano (*trachinotus carolinus*), and the black margate (*Anisotremus surinamensis*), use their pharyngeal jaws to crush the shells of snails and other mollusks in order to consume them (Lauder 1983; Grubich 2003; Gidmark et al. 2015). Others, such as the striped burrfish (*Chilomycterus schoepfi*), use their fused oral teeth to manipulate and crush shells (Winterbottom 1974; Ralston and Wainwright 1997). The biomechanical constraints of crushing hard shells is well documented in fish. For example, body mass (g), bite force (N), and pharyngeal jaw gape size are understood to limit the upper size of prey in the Caribbean hogfish (*Lachnolaimus maximus*), where larger fish generally produce both larger gapes and increased crushing force, allowing them to crush larger or thicker shells (Wainwright 1987, 1991). Similarly, the upper prey size consumed by black carp is limited by 1) the amount of force produced by its pharyngeal jaw closing muscle (*medial levator arcus branchialis V*) (Gidmark et al. 2013) and 2) the size of the pharyngeal jaw gape (Gidmark et al. 2015).

An alternative and much rarer method of consuming hard-shelled prey, primarily documented in cichlids endemic to Lake Malawi (*Metriaclima lanisticola*), Lake Victoria (*Hapochromis. xenognathus, H. sauvagei* and *Macropleurodus bicolor*), and Lake Edward *(H. concilians sp. nov., H. erutus sp. nov.* and *H. planus sp. nov*), is to extract the soft tissue of the gastropod from its shell via wrenching or shaking, known as ‘oral shelling’ (Slootweg 1987; Madsen et al. 2010; Lundeba et al. 2011; Vranken et al. 2019). It is typically thought that oral shelling is a way to circumvent the force and pharyngeal gape size requirements for consuming large mollusks because oral shelling does not require a fish to break a mollusk’s shell; however, very few studies have investigated oral shelling in general (but see: Slootweg 1987; De Visser and Barel 1996) nor have they investigated adaptations for oral shelling.

One possibility may be that fish use morphological adaptations to create a mechanical advantage during oral shelling. For example, one hypothesis is that the fleshy snout of *Labeotropheus* cichlids is used as a fulcrum, allowing fish to more easily crop algae from rocks versus the bite-and-twist method observed in other cichlid species (Konings 2007; Conith et al. 2018), and specifically that increased snout depth may help create this mechanical advantage (Conith et al. 2019). A similar method may be used during oral shelling to amplify force while removing snails from their shells. Thus, we predicted that larger nasal fulcrums should provide greater mechanical advantage for successfully oral shelling larger prey.

The durophagous pupfish (*Cyprinodon brontotheroides*) is an excellent species for testing whether a novel morphological trait provides a mechanical advantage for oral shelling. Durophagous pupfish are found within an adaptive radiation of pupfish endemic to the hypersaline lakes of San Salvador Island, Bahamas, which also includes a generalist pupfish (*C. variegatus*) and a scale-eating pupfish (*C. desquamator;* Martin and Wainwright 2011, 2013*a*). Geological evidence suggests that the hypersaline lakes of San Salvador Island, and thus the radiation itself, are less than 10,000 years old (Hagey and Mylroie 1995; Martin and Wainwright 2013*b*, 2013*a*). Phylogenetic evidence also indicates that: 1) generalist pupfish found outside San Salvador Island are outgroups to the entire San Salvador clade, and 2) that durophagous pupfish cluster near generalists from the same lake populations, indicating that there is extensive admixture between these young species (Martin and Feinstein 2014; Martin 2016; Lencer et al. 2017; Richards and Martin 2017). Gut content analyses indicated that durophagous pupfish consume approximately 5.5 times the number of mollusks and crustaceans (specifically ostracods) as generalists and fewer shells, suggesting that durophagous pupfish may be orally shelling their prey (Martin and Wainwright 2013*b*). In addition to their dietary specialization, durophagous pupfish also possess a novel nasal protrusion not observed in other pupfish species (Martin and Wainwright 2013*a*). This nasal protrusion is an expansion of the maxilla, and extends rostrally over the upper jaws (Hernandez et al. 2018). It is plausible that this nasal protrusion is an adaptation for oral shelling used by the durophage as a fulcrum.

We investigated oral-shelling behavior in the laboratory and tested if the nasal protrusion of durophagous pupfish is an adaptation for oral shelling. We measured snail consumption across 6 groups in the laboratory: outgroup generalists, generalists from San Salvador Island, scale-eaters, durophages, and F1 and F2 durophage hybrids (produced by crossing purebred durophages and generalists in the lab). If the novel nasal protrusion is adapted for oral shelling, we expected that durophages would consume significantly more snails than generalists and scale-eaters. We also expected that F1 hybrids would show intermediate snail consumption between the parental species and that F2 hybrids would show greater variation in snail consumption compared to parental species. To directly tie nasal protrusion size to snail-shelling performance, we also investigated whether individuals with larger noses could consume larger snails in lab-reared populations of both durophages and F2 hybrids. Ultimately, we found that, contrary to our predictions, purebred durophages, F1, and F2 hybrids all shelled significantly more snails than other pupfish species and we did not find evidence that larger nasal protrusion allowed durophages to consume larger snails. Instead, we discuss alternative explanations for the novel nasal protrusion such a putative function in foraging efficiency, sexual selection, olfaction, or increased area for superficial neuromasts.

## Methods

### Collection and Care

During the summer of 2017, we used seine nets to collect generalist, durophage, and scale-eater pupfishes from Crescent Pond, Little Lake, Osprey Lake, and Oyster Pond (San Salvador Island, Bahamas). We transported fish back to the University of North Carolina, Chapel Hill, where they were maintained in mixed-sex stock tanks (37-75 l) in approximately 26° C water at approximately 5-10 ppt salinity (Instant Ocean salt mix). In the lab, we produced F1 and F2 hybrid offspring using snail-eater and generalist parents. Wild caught individuals were also allowed to breed and produced F1-F3 purebred offspring. Hybrid and purebred offspring were used in our feeding assays. We fed all fish a diet of commercial pellet foods, frozen bloodworms, and mysis shrimp daily.

We also maintained a colony of freshwater sinistral snails (*Physella sp.*). We kept snails in a 7 liter stock tank containing the same water used in pupfish tanks. All snails were acclimated to 5-10 ppt salinity for at least 48 hours before being used in a feeding trial. We fed snails a diet of bloodworms every 48 hours. We ran multiple control trials without fish alongside feeding trials to track natural snail mortality rates.

### Morphological Measurements

We measured standard length of each fish by measuring the distance from the tip of the upper jaw to the posterior end of the hypural plate. We also measured nasal protrusion size for a subset of fish (9 generalists, 50 durophages, 17 F1 hybrids, and 62 F2 hybrids) using image processing software (Schindelin et al. 2012). Scale-eating pupfish do not exhibit even marginal nasal protrusion, and therefore we did not include them in this analysis. We measured fish nasal protrusion size by drawing a tangent line aligning the most anterior dorsal point of the premaxilla with the neurocranium and measuring a perpendicular line at the deepest part of the nasal region (Figure 1C).

**Figure 1.**
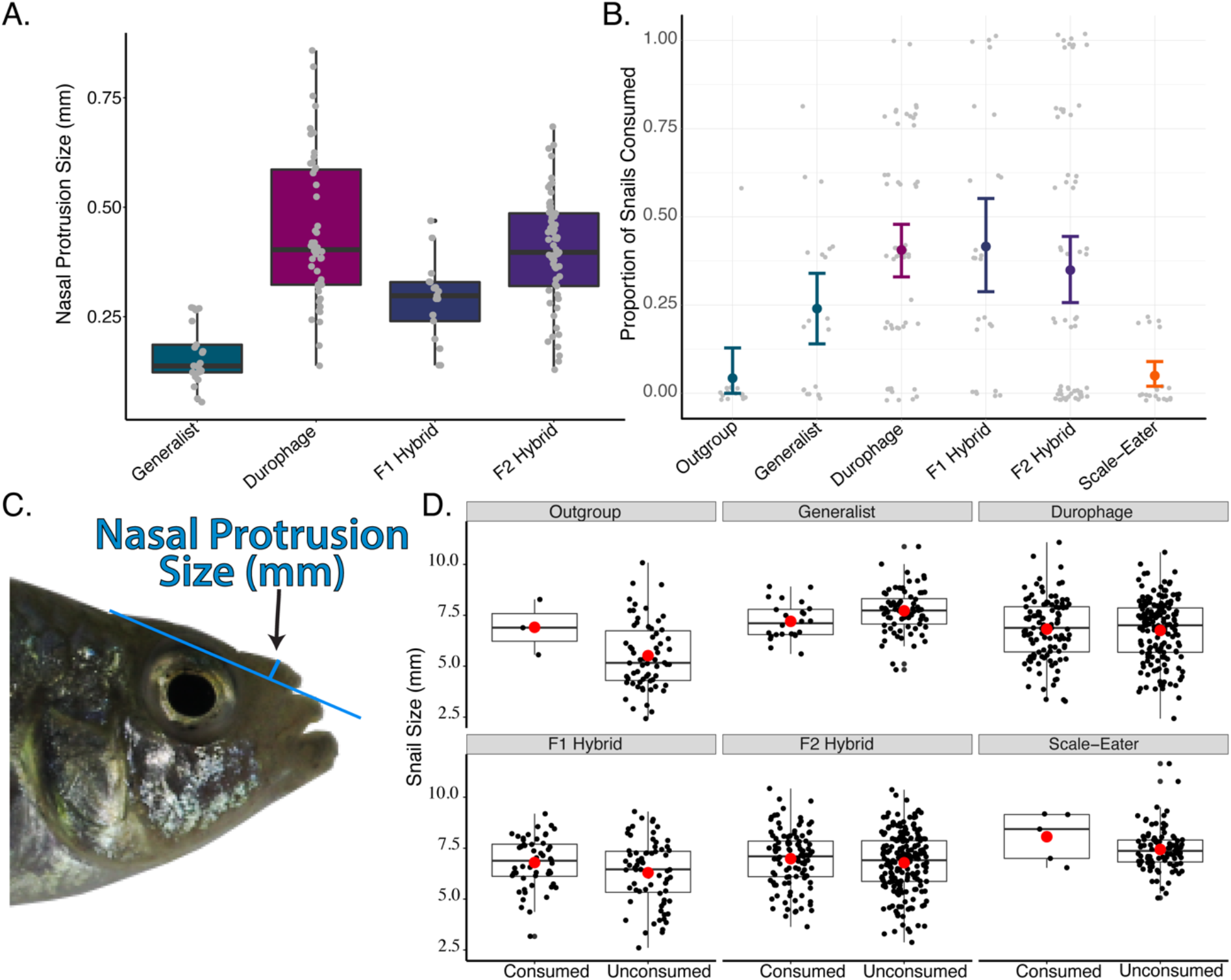
Snail consumption, nasal protrusion size, and snail size by species. A) Variation in nasal protrusion size across pupfish groups. Grey dots represent individual fish. B) Proportion of snails consumed across six groups of pupfish. Colored dots represent mean proportion, and error bars represent 95% confidence intervals (bootstrapping: 1,000 iterations). C) Visualization of how nasal protrusion size was measured (pictured: durophagous pupfish). D) Visualization of the size of consumed and unconsumed snails for each species. Black dots represent individual snails and red dots represent the mean snail size.

### Feeding Assay

We quantified the number of snails consumed by all three species of pupfish and hybrids using feeding assays. Prior to a feeding assay, fish were removed from stock tanks and isolated in 2L trial tanks which contained one synthetic yarn mop to provide cover for the fish. We allowed fish to acclimate in trial tanks for at least 12 hours before the start of a feeding assay. After the acclimation time, we haphazardly chose 5 snails from our snail stock tank and added them to each feeding assay tank. We added one bloodworm to each tank to ensure that even fish which did not consume any snails had an adequate diet. Fish were allowed to feed freely on snails for 48 hours with no additional food source. At the end of the 48-hour assay period fish were removed from trial tanks, photographed, and placed back into mixed-sex stock tanks. We then recorded the number of snails that were consumed (empty shells remaining) and unconsumed. Finally, we measured the size of each snail shell from the anterior tip of the shell’s aperture to farthest tip of the spire (mm) using digital calipers and image processing software. In total, we measured feeding success for 13 outgroup generalists, 20 generalists, 55 durophages, 20 scale-eaters, 25 F1 hybrids, and 63 F2 hybrids. Out of the 196 trials, only 11 finished the trial period with four snail shells instead of the given five, suggesting that at most 3.5% of snail consumption involved also eating the shell.

### Data Processing

#### No differences between fully consumed and partially consumed snails

We noticed that a portion of the snails were only partially consumed (i.e. part of the snail tissue remained in the shell versus a completely empty shell after 48 hours) and therefore used a generalized linear mixed model (GLMM) with a binomial response distribution to determine if partially consumed snails should be analyzed separately from fully consumed snails. We included 1) whether snails were fully or partially consumed as the response variable (binomial data), 2) species designation as a fixed effect, 3) population and fish ID as random effects, and 4) log standard length as a covariate. We found that the pattern of partially and fully consumed snails did not vary across species (*χ*^2^= 2.73, *df*=5, *P*=0.74), and therefore included all partially consumed snails in the general “consumed” category for the remainder of our analyses.

#### Statistical Analysis

We used a linear mixed model to investigate the relationship between nasal protrusion distance and species. For this analysis we used a subset of our data which includes: 9 generalists, 50 durophages, 17 F1 hybrids, and 62 F2 hybrids. Our model included 1) log nasal protrusion size as the response variable, 2) species designation, log standard length, and their interaction as fixed effects, and 3) population as a random effect. We also used Tukey’s HSD to make *post hoc* comparisons across species.

We used a GLMM with a negative binomial distribution to explore whether the number of snails consumed varied between species. We included 1) whether snails were consumed or unconsumed as the response variable (binomial data), 2) species designation as a fixed effect, 3) population and fish ID as random effects, and 4) log standard length as a covariate. We made additional *post hoc* comparisons between groups using Tukey’s HSD.

We used a linear mixed model to determine if the size of snails varied by whether they were consumed or unconsumed and whether that varied between species. We included 1) snail size (mm) as the response variable, 2) whether snails were consumed or unconsumed, species designation, and their interaction as fixed effects, 3) population and fish ID as random effects, and log standard length as a covariate. We made additional *post hoc* comparisons between groups using contrasts and an FDR correction.

Finally, we investigated if nasal protrusion distance affected the maximum size snail an individual could consume as an estimate of snail-shelling performance. For this analysis we only considered purebred durophages and F2 hybrids (separately) as they had the largest observed variance in nasal protrusion size and only included individuals that consumed at least one snail during the feeding trial. For each group, we used a linear model with 1) the size of the largest consumed snail for each individual as the response variable, 2) log nasal protrusion size, log standard size, and their interaction as fixed effects, and 3) the residuals from a linear model investigating the relationship between snail size and nasal protrusion size as a covariate. We included this additional covariate because we found a strong positive relationship between mean snail size provided during trials and nasal protrusion in both purebred durophages (LM: *P*=1.72 × 10^−9^, adjusted R^2^ =0.14) and F2 hybrids (LM: *P*=5.58 × 10^−10^, adjusted R^2^ =0.12), and wanted to account for this variation in the model (Figure S2). This variation reflected our attempt to provide some larger snails in trials with larger fish to better assess performance. We additionally included the random effect of population in our durophage model.

#### Ethical Statement

This study was conducted with the approval of the Animal Care and Use Committee of the University of North Carolina, Chapel Hill, NC (protocol# 15–179.0). All wild fish were collected with a research and export permit from the Bahamas BEST commission, renewed annually since 2011.

## Results

### Nasal protrusion size does not vary between purebred durophages and hybrids

Our linear mixed model indicated that nasal protrusion size is significantly associated with log standard length (*χ*^2^= 27.63, *df*=1, *P*=1.47×10^−7^), but that this relationship does not vary between purebred and hybrid durophages (*χ*^2^= 3.22, *df*=3, *P* = 0.36; Figure 1A & S1). *Post hoc* analysis indicated that generalists had smaller noses than durophages (*P* < 0.0001) and F1 hybrids (*P* = 0.016).

### Purebred durophages and their hybrids consume the most snails

We found that species designation was a significant predictor for the number of snails an individual consumed (GLMM; *χ*^2^= 35.61, *df*=5, *P*= 1.129×10^−6^). Specifically, we found that durophages, F1 hybrids, and F2 hybrids consumed more snails than the generalist outgroup population (Lake Cunningham, New Providence Island, Bahamas) and scale-eating pupfish (Figure 1B). Durophages, F1 hybrids, and F2 hybrids also consumed twice as many snails as generalists, however this difference was not significant.

### Consumed snails were larger than unconsumed snails

In general, we found that the size of snails varied 1) by whether they were consumed (*χ*^2^= 4.002, *df*=1, *P*=0.045), and 2) across species (*χ*^2^= 24.79, *df*=5, *P*=0.00015; Figure S1). Specifically, we found that consumed snails were on average 0.12 mm larger in diameter than unconsumed snails (*P*=0.046). Generalists and scale-eaters received snails that were approximately 17% larger than other groups (generalists: *P*=0.016; scale-eaters: *P*=0.02). Although this was unintentional due to the available size distributions of snails in our colony over the ten month course of the feeding trails, we believe that it did not introduce a significant bias because 1) larger snails were more likely to be consumed (in fact there was only an 8% difference between the mean size of snail given to generalists and scale-eaters *vs* the mean size of consumed snails) and 2) generalists and scale-eaters were excluded from analyses which examined how nasal protrusion affected a fish’s ability to consume snails.

### Nasal protrusion size did not significantly increase the maximum snail size consumed

We found no effect of log nasal protrusion size, log standard length, or their interaction on the size of the largest consumed snail for either durophages (*P_log(nasalprotrusionsize)_*=0.49, *P*_*log(standardlength)*_=0.61, *P*_*interaction*_=0.56) or F2 hybrids (*P*_*log(nasalprotrusionsize)*_=0.83, *P*_*log(standardlength)*_=0.66, *P*_*interaction*_=0.91; Figure 2).

**Figure 2.**
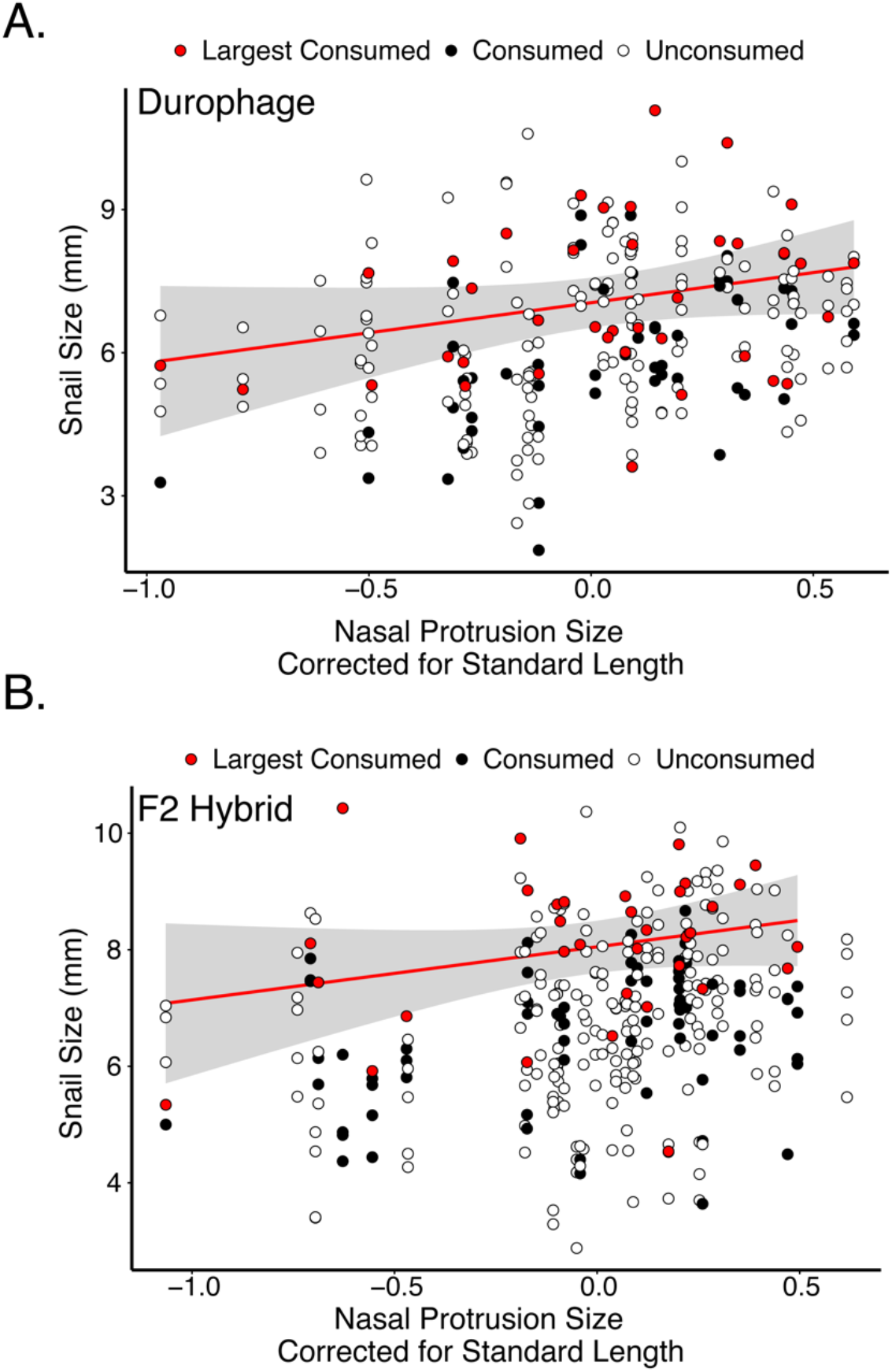
The maximum prey size a pupfish can consume was not affected by nasal protrusion size. The X-axis shows nasal protrusion size corrected for standard length while the Y-axis shows snail size (mm). Red dots show the size of largest consumed snail from each trial, the red line represents the linear model describing the relationship between nasal protrusion size and the largest consumed snails, and the grey area represents 95% CI. Closed circles show the size of other snails that were consumed during trials; open circles show the size of unconsumed snails.

## Discussion

We present the first strong evidence in any cyprinodontiform fish that the durophagous pupfish is an oral sheller, shaking snails free from their shells rather than crushing or ingesting the whole shell. This is consistent with their notably non-molariform pharyngeal jaws relative to generalists and snail-crushing species (Figure 3). We then tested the hypothesis that the durophagous pupfish’s novel nasal protrusion is an adaptation for removing snails from their shells, potentially functioning as a fulcrum. We predicted that durophagous pupfish would 1) consume more snails than other groups, and 2) consume larger snails than other groups. We found that both durophages and their F1 and F2 hybrid offspring consumed the most snails compared to other groups (Figure 1B), indicating that any substantial amount of durophagous genetic ancestry increases the number of snails consumed over a 48-hour feeding trial. However, contrary to our expectations, we found no significant evidence that larger nasal protrusions within hybrid or parental durophagous pupfish populations enabled the fish to consume larger snails (Figure 2).

**Figure 3.**
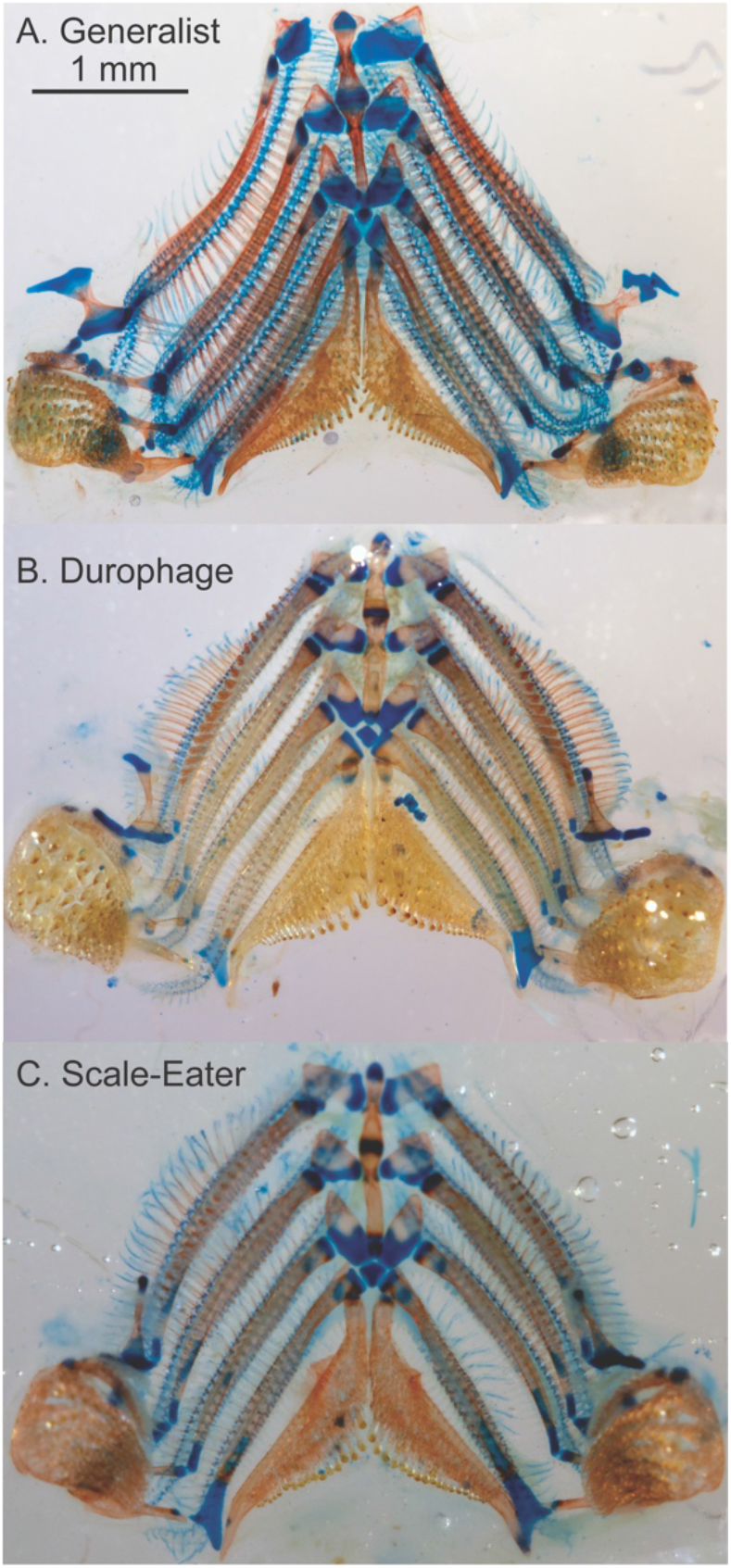
Branchial skeleton and pharyngeal teeth of all three San Salvador Island species. Image of the dissected branchial skeleton and pharyngeal jaws of A) generalist, B) durophage, and C) scale-eater pupfish. Scale (1mm) is shown in Figure A and is consistent across all three photos. From these three individuals, the representative snail-eater has lower pharyngeal teeth that are 50% longer and 75% wider than the generalist or scale-eating individuals.

### Durophages have a stronger behavioral preference for snails compared to other species

One explanation for the observed pattern is that durophagous pupfish have a stronger preference for snails which is independent from their novel nasal protrusion. We see some support for this within our data. Generalist pupfish from San Salvador Island consumed significantly more snails than generalists found outside of the radiation on New Providence Island, and even consumed statistically similar amounts of snails as purebred durophages despite having much smaller nasal protrusions (Figure 1A&B). It could be that extensive geneflow between generalists and durophages on San Salvador Island spread alleles for snail-eating preference throughout both pupfish species (Martin and Feinstein 2014). Alternatively, the common ancestor of durophages and generalists may have had a strong preference for snails (Martin and Feinstein 2014; Richards and Martin 2017). The increased aggression of both male and female durophages toward conspecifics by potentially alternate genetic pathways to scale-eaters, as shown in a recent study (St. John et al. 2019), could also be associated with their stronger preference for aggressively attacking snails to flip them over before gripping the body of the snail in their oral jaws and shaking them free from their shells (Supplemental Video 1).

Liem’s hypothesis and subsequent work has long supported the idea that morphological specialization need not coincide with trophic specialization, or *vice versa*. For example, *Tropheops tropheops* and *Metriaclima zebra*, two cichlids from Lake Malawi that are morphologically specialized for scraping algae often fill a generalist ecological niche, consuming zooplankton, benthic invertebrates, and phytoplankton (Liem 1978, 1980; McKaye and Marsh 1983), particularly during periods of resource abundance (Martin and Genner 2009). An analogous argument can be made for individual dietary specialization within a population (Bolnick et al. 2003). For example, Werner and Sherry (1987) found that individual Cocos Island finches specialize on a wide variety of taxa including crustacea, nectar, fruit, seeds, mollusks, and lizards, and that individual dietary specialization was most likely driven by behavioral differences. Similarly, increased levels of individual specialization in sticklebacks are driven by shifts in forager density or intraspecific competition (Svanbäck and Bolnick 2005, 2007; Araújo et al. 2008). Thus, individual specialization is often driven entirely by differences in behavior, feeding preference, or other external factors and can be divorced from adaptive differences in morphology (Werner and Sherry 1987).

### Alternative functions of the novel nasal protrusion

We investigated whether an increase in nasal protrusion size affected the maximum size snail an individual could consume (Figure 2). However, it could be that the novel nasal protrusion is related to feeding efficiency, e.g. in handling time per snail, or is a sensory organ used for locating snails more efficiently with potentially increased numbers of superficial neuromasts (Shibuya et al. 2019). There are several examples of nasal protrusions that are used for this purpose. The unique rostrums of paddlefish (Polydontidae), sturgeon (Acipenseridae), and sawfish (Pristidae) are all used as sensory organs, containing electroreceptors, lateral line canals, and even barbels for detecting prey items (Miller 2006; Wueringer et al. 2012). The novel nasal protrusion of the durophagous pupfish may also be a sensory organ, however, whether the nasal protrusion has an increased number of superficial neuromasts is still unknown.

Alternatively, the novel nasal protrusion may allow durophagous pupfish to orally shell snails more quickly, increasing their feeding efficiency. For example, Schluter (1993) documented that benthic sticklebacks with deep bodies, large mouths, and few, short gill rakers were more efficient at consuming benthic prey items, while limnetic species of stickleback, with slender bodies, small mouths, and many, long gill rakers, were more efficient at consuming limnetic prey items. Interestingly, Schluter (1993, 1995) also found that F1 hybrids had decreased efficiency feeding on both limnetic and benthic prey items which was primarily due to their intermediate phenotypes and suggested that reduced fitness in hybrids helps maintain species boundaries between benthic and limnetic species. It could be that the durophage F1 and F2 hybrids have similar preferences for gastropods, but cannot consume snails as efficiently due to their intermediate phenotype. However, we found no strong evidence suggesting that the nasal protrusion is adapted for oral shelling (Figure 2). Future work should investigate other traits that may be adaptive for oral shelling such as the strength of the dorsal head of the maxilla which comprises the skeletal basis of the novel nasal protrusion, structural differences in the mandibular symphysis, coronoid process, or the articular bones which may all provide additional strength or stabilization during biting, or tooth variation in the durophage pharyngeal jaws (Fig. 3). Indeed, there is subtle variation apparent in the pharyngeal teeth and jaws of durophages compared to other pupfish species (Figure 3) which has not been previously reported, suggesting that pharyngeal jaws may be adapted for processing hard-shelled prey.

### The novel nasal protrusion may be a sexually selected trait

Finally, the novel nasal protrusion may be unrelated to oral shelling and instead may be used in species recognition or mate preference functions. Exaggerated traits, like the novel nasal protrusion in durophage pupfish, commonly arise via sexual selection. For example, forceps size in earwigs (Simmons and Tomkins 1996), major claw size in fiddler crabs (Rosenberg 2002), and the size of the sword tail ornament present in swordtail fish (Rosenthal and Evans 1998) are all thought to be sexually selected traits. Two commonly invoked hallmarks of a sexually selected trait are 1) allometric scaling compared to body size and 2) that the trait is sexually dimorphic (Kodric-Brown and Brown 1984; Kodric-Brown et al. 2006; Shingleton and Frankino 2013). In pupfish, there is a weak positive relationship between standard length and nasal protrusion size observed for generalists (Figure S1A, generalist_slope_= 0.35). Generalist pupfish mostly likely resemble the most recent common ancestor for the radiation, making the observed slope a good null expectation for how nasal protrusion size should scale with body size in pupfish. In durophages, we observe much stronger positive allometry of the nasal protrusion (Figure S1B, durophage_slope_= 0.93), in which large durophage individuals have nasal protrusion sizes more than twice as large as those in large generalists. However, we found no significant difference in nasal protrusion size between male and female durophages when accounting for these size differences (LM, *P*=0.96).

## Conclusion

In conclusion, we did not find evidence to support that the novel nasal protrusion observed in durophagous pupfish is adapted for consuming large snails. Instead, we found that purebred durophages and their F1 and F2 hybrids have stronger preferences for consuming snails than other species. We suggest that the novel nasal protrusion may be adapted for other aspects of oral shelling such as feeding efficiency, or that variation in other traits, such as the pharyngeal jaws (Figure 3), may play a larger role in oral shelling. Alternatively, this may be an example of trophic specialization due to behavioral specialization (i.e. feeding preference).

